# Giardia colonizes and encysts in high density foci in the murine small intestine

**DOI:** 10.1101/080226

**Authors:** NR Barash, C Nosala, JK Pham, SG Mclnally, S Gourguechon, B McCarthy-Sinclair, SC Dawson

**Affiliations:** Department of Microbiology and Molecular Genetics, One Shields Avenue, UC Davis, Davis, CA 95616; Department of Molecular and Cell Biology, 345 LSA Bldg. UC Berkeley, Berkeley, CA 94720

## Abstract

*Giardia* is a highly prevalent, yet understudied protistan parasite causing diarrheal disease worldwide. Hosts ingest *Giardia* cysts from contaminated sources. In the gastrointestinal tract, cysts excyst to become motile trophozoites, colonizing and attaching to the gut epithelium. Trophozoites later differentiate into infectious cysts that are excreted and contaminate the environment. Due to the limited accessibility of the gut, the temporospatial dynamics of giardiasis in the host is largely inferred from laboratory culture and thus may not mirror *Giardia* physiology in the host. Here we have developed bioluminescent imaging (BLI) to directly interrogate and quantify the *in vivo* temporospatial dynamics of giardiasis, thereby providing an improved murine model to evaluate anti*-Giardia* drugs. Using BLI, we determined that parasites primarily colonize the proximal small intestine non-uniformly in high-density foci. By imaging encystation-specific bioreporters, we show that encystation initiates shortly after inoculation and continues throughout the entire duration of infection. Encystation also initiates in high-density foci in the proximal small intestine, and high-density laboratory cultures of parasites are also stimulated to encyst. This work overturns the assumption that parasites encyst later during infection as they are dislodged and travel through the colon. We suggest that these high-density regions of parasite colonization likely result in localized pathology to the epithelium, and encystation occurs when trophozoites reach a threshold density due to local nutrient depletion. This more accurate visualization of giardiasis redefines the dynamics of *in vivo Giardia* life cycle, paving the way for future mechanistic studies of density-dependent parasitic processes in the host.

**Significance:** *Giardia* is a single-celled parasite causing both acute and chronic diarrheal disease in over one billion people worldwide. Due to limited access to the site of infection in the gastrointestinal tract, our understanding of the dynamics of *Giardia* infections in the host has remained limited, and largely inferred from laboratory culture. To better understand giardiasis in the host, we developed imaging methods to quantify *Giardia* expressing bioluminescent physiological reporters in live mice. We discovered that parasites primarily colonize and encyst in the proximal small intestine in discrete, high-density foci. Furthermore, this work provides evidence of a parasite density-based threshold for the differentiation of *Giardia* into cysts in the host. These findings overturn existing paradigms of giardiasis infection dynamics in the host.

## INTRODUCTION

*Giardia lamblia* is a unicellular protistan parasite causing acute and chronic diarrheal disease in over one billion people worldwide, primarily in developing countries with inadequate sanitation and water treatment (1, 2). Giardiasis is a serious disease of children, who may experience substantial morbidity including diarrhea, malnutrition, wasting, and developmental delay (3-5). In the United States, giardiasis is the most frequently diagnosed cause of water-borne diarrheal disease, and commonly affects travelers and immunosuppressed individuals (6). Trophozoites are not invasive and giardiasis does not produce a florid inflammatory response; however, giardiasis is associated with villus shortening, enterocyte apoptosis, hypermobility, and intestinal barrier dysfunction (7). The estimated failure rates of up to 20% for standard drug treatments such as metronidazole (8) and growing evidence of drug resistance in *Giardia* (9-11), underscore the need for new therapeutic treatments of this widespread and neglected diarrheal disease.

Motile *Giardia* trophozoites colonize and proliferate in the small intestine (12), attaching to the intestinal villi to resist peristalsis using a complex microtubule structure termed the ventral disc (13,14). In the gut, trophozoites differentiate into infectious cysts that are eventually excreted and can contaminate water sources in the environment (6,15). Disseminated cysts are ingested from contaminated water, and then excyst into trophozoites after passage through the stomach, completing their life cycle in the host gastrointestinal tract. Trophozoites are proposed to colonize the acidic, cholesterol-rich jejunum and then initiate encystation when peristalsis sweeps them to the alkaline, cholesterol-depleted distal intestine (16-18). Encystation is thus believed to be triggered via cues from a specific gastrointestinal anatomical site (17,19), and can be induced *in vitro* by lowering pH and cholesterol, and increasing bile and lactic acid in the medium (16, 20). However, cysts produced *in vitro* are less robust at establishing infections in animal models than cysts harvested directly from feces, implying that additional host factors are required for infectious cyst production (17).

As differentiation of the trophozoite into the infectious cyst is a critical aspect of *Giardia’s* pathogenesis (21), determining the extent of *in vivo* parasite differentiation to cysts and subsequent cyst dissemination is key to understanding *in vivo* host-parasite interactions (22-24). Despite decades of study, there remains a lack of understanding of the *in vivo* host-parasite infection dynamics underlying the extent and progression of acute and chronic giardiasis (5, 25-27). Animal models of giardiasis include adult (28, 29) or suckling mice (30) or adult gerbils (31) infected with either *Giardia lamblia or G. muris* isolates (32). The adult mouse model of giardiasis is commonly used to evaluate anti-giardial drugs (33). Observation and quantification of *in vivo* parasite physiology and differentiation has primarily been indirect, however, and *in vivo* physiological studies are have been contingent on isolating parasites from sites of infections in animal models post-sacrifice.

Due to the limited accessibility of the gastrointestinal tract (16-18), our understanding of the temporal and spatial infection dynamics of giardiasis is largely inferred from laboratory culture rather than *in vivo* models of the disease (16,18). While *in vitro* studies have established that the initiation of encystation is transcriptionally controlled (9-11), understanding the complex spatiotemporal dynamics of the parasite life cycle and interactions with the host remains challenging. *In vitro* models of giardiasis are not adequate proxies for infection within the host as they do not accurately mirror *in vivo* parasite physiology, and *in vitro* studies rarely have been confirmed through analogous *in vivo* studies of parasite physiology. Thus, *in vivo* models are necessary to understand parasite infection dynamics in the host and to evaluate new anti-giardial drugs.

To assess parasite colonization and differentiation dynamics in the host, we developed bioluminescent imaging methods (BLI) allowing us to directly quantify and image temporal and spatial dynamics of giardiasis. Specifically, we have infected mice with trophozoites expressing firefly luciferase under the control of constitutive or encystation-specific (34-38) promoters. BLI enables sensitive quantification and live reporting of transcriptional activity of protein fusions (39-41), and has been used previously to monitor parasitic infection dynamics during malaria, leishmaniasis, trypanosomiasis, and toxoplasmosis (42-44), as well as bacterial colonization of the intestine (34). Non-invasive imaging of bioluminescent *Giardia* parasites permits unprecedented access to real-time host-parasite interactions, allowing us to revise decades-old assumptions about *Giardia* infection and encystation dynamics in living hosts. Importantly, we demonstrate that metabolically active parasites primarily colonize the proximal small intestine, rather than colonizing the mid-jejunum as has been previously reported in adult mice (28) and immunodeficient mice (17), or inferred from the identification of cysts in distal anatomical sites of the gastrointestinal tract (e.g. ileum and colon) (20, 23). Further, we improve our understanding of the *in vivo Giardia* life cycle, demonstrating that encystation is initiated early in the course of infection, peaks within first week, and is correlated with the highest parasite density during infection. Quantification of encystation-specific vesicles (ESVs) in trophozoites colonizing the proximal small intestine confirms the observations made using BLI. Lastly, we demonstrate that local parasite density in the host contributes to the induction of encystation-specific transcription *in vitro* and may contribute to the early *in vivo* differentiation of parasites in mice. Using BLI to evaluate *in vivo* giardiasis permits precise longitudinal and spatial monitoring of the dynamics of infection, and provides an improved method to evaluate anti-giardial drugs in a relevant animal model of giardiasis.

## RESULTS

### Visualizing and quantifying Giardia infection dynamics using non-invasive bioluminescent imaging

To confirm that the promoter-firefly luciferase (FLuc) fusions are stably integrated and that the bioluminescence is not lost in the absence of antibiotic selection (Supplemental Figure SI), we used *in vitro* bioluminescence assays to monitor luciferase activity after the removal of antibiotic selection (Supplemental Figure S2B). Both *P_GDH_-FLuc* and *P_CWP1_-FLuc* strains maintained a consistent bioluminescence for at least three weeks under normal growth conditions (Supplemental Figure S2B). Luciferase catalyzes the production of light in the presence of luciferin substrate. While oxygen is required for light production, the colon has sufficient oxygen for detectible light output (45). Because *Giardia* trophozoites proliferate in the low oxygen gut lumen, we tested D-luciferin delivery both orally (by gavage) and systemically (by intraperitoneal injection) to determine the delivery method that produced the optimal bioluminescent signal (Supplemental Figure S3A). Intraperitoneal injection caused animals less stress and produced stable bioluminescence from bioreporter strains for over 30 minutes after injection (Supplemental Figure S3B). Uninfected mice or mice infected with a non-luminescent strain of *Giardia* had negligible background signal (Figure 1A).

To query the temporal sequence of both *in vivo* colonization and encystation, we infected a cohort of mice with one million *P_GDH_-FLuc* trophozoites and quantified the bioluminescent signal over a 14-day time course (Figure 1). At day 7, we observed significant bioluminescence as compared to uninfected animals (ratio paired T test, p<0.0067). Individual mice showed variation in the degree of bioluminescent signal (Figure 1A) and some animals exhibited signal periodicity; one representative individual (Figure IB) showed bioluminescence peaks at day 4, day 10 and day 12 post-infection. Maximum bioluminescence occurred between day 4 and day 9 for all animals infected with the *P_GDH_-FLuc* strain (n=20 over two experiments, Figure 1C). To ensure the bioluminescent signal was attributable to metabolically active parasites, we also treated mice infected with the *P_GDH_-FLuc* strain with 50 mg/kg metronidazole by oral gavage. After two days of treatment, the bioluminescent signal had decreased to the same level as non-infected animals (Figure ID).

**Figure 1.**
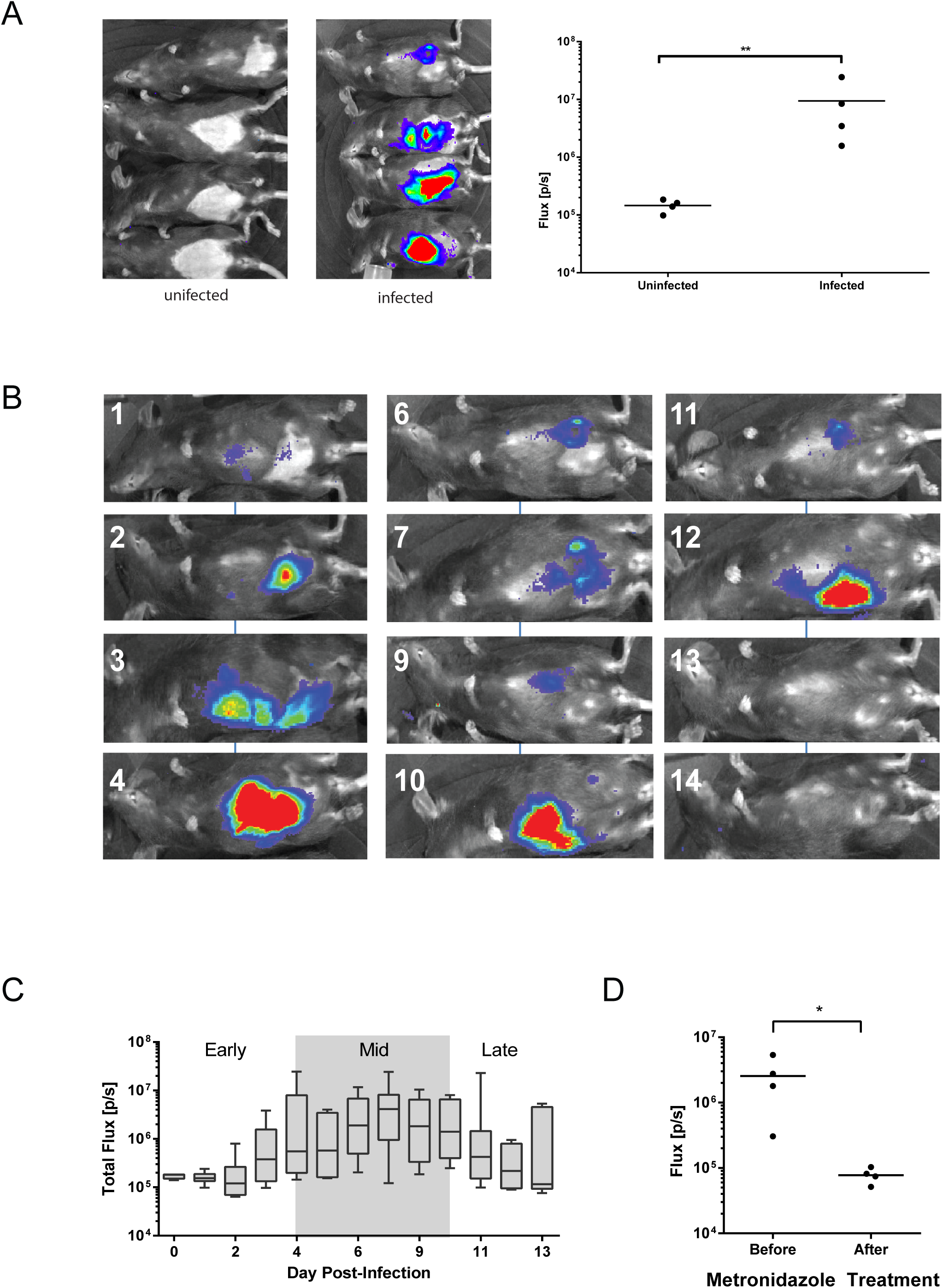
Bioluminescent imaging allows non-invasive quantification of Giardia infection dynamics in individual cohorts of mice. In panel A, a cohort of four of mice were inoculated with the *P_GDH_-FLuc* strain and bioluminescence at seven days post-inoculation was compared to the background signal from uninfected animals (images left). *P_GDH_-FLuc* bioluminescence intensity is indicated in the image overlays, with the highest signal intensity shown in red and the lowest in blue. Bioluminescence flux (photons/second) is compared between the uninfected and infected group, with the double asterisk indicating significance as assessed by the ratio paired T test (p<0.0067). In B, cyclic variability of the *P_GDH_-FLuc* bioluminescence is shown in the same animal imaged non-invasively each day for fourteen days post infection. Infection dynamics in two experiments with 8 and 12 mice as quantified by *P_GDH_-FLuc* bioluminescence is summarized in panel C. The average bioluminescence measured for the cohort each day is shown. The shaded box indicates the maximal bioluminescent signal, or peak infection range. The average time to maximal bioluminescence after infection with the *P_GDH_-FLuc* was 6.6 days. Lastly, in D, a cohort of four mice was imaged and flux (p/s) was quantified five days after infection with the *P_GDH_-FLuc* strain (Before). The same four mice were imaged and flux was quantified two days after treatment with 50 mg/kg metronidazole by oral gavage (After).

### Quantifying the spatial variation of giardiasis using ex vivo imaging of the gastrointestinal tract

To assess spatial infection dynamics and to correlate non-invasive imaging with *ex vivo* imaging of excised intestine, we inoculated twenty-one mice with one million *P_GDH_-FLuc* trophozoites. On days 1, 3, 5, 7, 9,11, and 13 post-infection, three or four animals were individually imaged. Animals were then sacrificed, and the gastrointestinal tracts were quickly excised and imaged *ex vivo* (Figure 2). We observed four major patterns of bioluminescence within the gastrointestinal tracts over the course of infection (representative patterns are shown in Figure 2A). The majority of bioluminescent signal occurred in the proximal small intestine as early as one day following oral gavage (Figure 2B), yet there was some spatial variability in the gastrointestinal parasite colonization pattern in the cohorts over the thirteen days. Further, we observed localized areas of maximal bioluminescent signal, or foci, within colonized regions of the gut (Figure 2A). These regions are upwards of 100 fold more bioluminescent than adjacent regions in the same anatomical section. In some animals, bioluminescence was present in the distal small intestine or diffuse throughout the small intestine. Less commonly observed was bioluminescence occurring primarily in the cecum or the large intestine. For all samples, BLI signal intensities of less than 1% of total maximal signal were seen within the stomach. The *in vivo* imaging signal intensities were directly comparable with the *ex vivo* imaging (Supplemental Figure S5).

**Figure 2.**
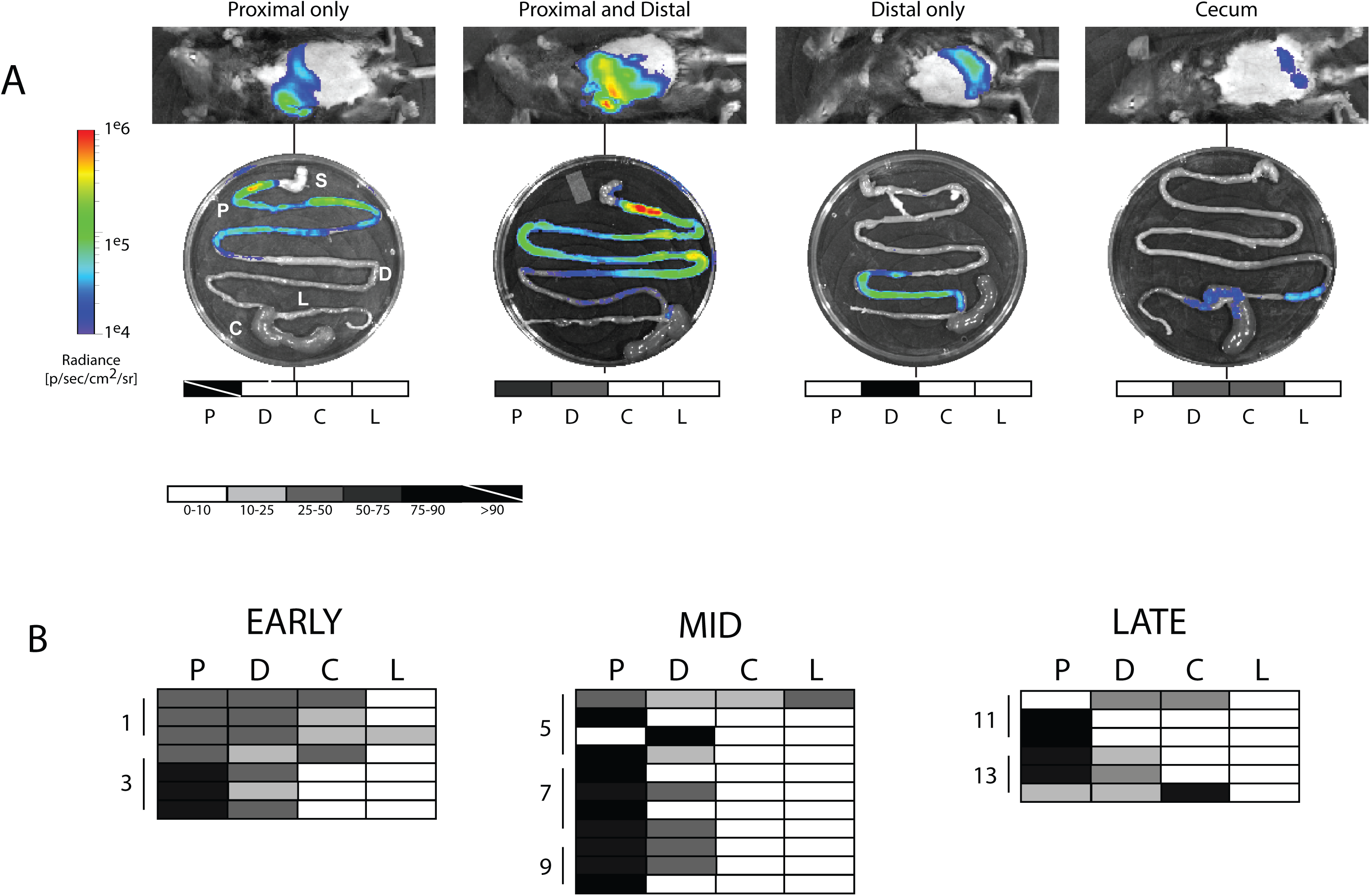
High-density foci of Giardia colonization are present in the proximal small intestine. Representative classes of *in vivo* and *ex vivo* bioluminescent images are shown for twenty-four mice infected with the *P_GDH_-FLuc* strain and sacrificed in cohorts of four on days 3, 5, 7, 9,11, and 13 post-infection (A). Photon flux or radiance (p/s/cm^2^/sr) for each intestinal segment is shown and has been normalized to the maximal *ex vivo* bioluminescence signal on the radiance scale, yielding the percent total signal per segment. These values are represented graphically on the gray scale maps below each *ex vivo* image (clear = 0-10% and black = 75-100%, with values between 10% and 75% indicated as shades of gray). A strike through indicates greater than 90% of maximal bioluminescent signal. In (B), quantitative bioluminescence imaging from infections is categorized and summarized by the region of the gastrointestinal tract (P = proximal small intestine, D = distal small intestine, C = cecum, and L = large intestine) for all animals in each phase of infection as **early** (days 0-3), **peak** (days 5-9), and **late** (days 11-13). The stomach (S= stomach) is shown for orientation but always lacks bioluminescence. Shading in each row of the charts indicates the variations in the maximal bioluminescence in each of the four regions in an individual animal.

Through a comparison of colonization patterns during early, mid, and late infections (Figure 2B), we found that early in infection, there was more diffuse small intestinal colonization, with 48% of the BLI signal from all animals localized to the proximal small intestine and nearly one third of signal from the distal small intestine. At maximal infection, the proximal small intestine was more strongly colonized than the distal, accounting for 71% of overall signal. Four of eleven mice (36%) had a proximal-only colonization pattern, with an average of 89% proximal signal amongst the individuals. Only one mouse had significant colonization of the distal intestine. Late in infection, higher BLI signal intensity was detected in the distal small intestine and cecum, although the proximal small intestine still accounted for 57% of overall signal. Early and late infections were characterized by a more diffuse pattern throughout the gastrointestinal tract, whereas during the maximal infection (mid infection) more parasites are concentrated in the proximal small intestine.

We next compared parasite quantification obtained using the BLI signal of the *P_GDH_-FLuc* bioreporter with other methods of parasite quantification. Following *ex vivo* imaging and quantification of bioluminescence, we quantified total parasites using qPCR of genomic DNA isolated from one-centimeter intestinal segments in regions of high and low bioluminescent signal (Supplemental Figure S4). We amplified the *Giardia* pyruvate ferredoxin oxidoreductase gene *(PFOR1)* and used the constitutively expressed murine *nidogen*-1 gene as an internal control to determine the contribution of murine DNA to total genomic DNA isolated from intestinal segments. A smaller difference in differential counts to threshold (ACT) between *nidogen* and *>PFOR*> indicated greater parasite as more murine DNA was present than *Giardia* DNA, while a larger difference in ΔCT indicated fewer parasites. We determined that there is a significant and linear association between bioluminescence intensity and infection density (Supplemental Figure S4, p < 0.0001).

### Encystation occurs early in infection in both the proximal and the distal small intestine

*Giardia* cysts consist of a partially divided trophozoite surrounded by a desiccation resistant cyst wall that is composed predominantly of leucine-rich cyst wall proteins (CWPs). CWPs are transported to the outer membrane by encystation-specific vesicles (ESVs) approximately six hours after transfer to *in vitro* encystation medium (23, 46). Cyst wall protein 1 (CWP1) expression is upregulated several hundred fold within seven hours after switching to *in vitro* encystation medium (23) (47). The bioluminescent signal from the *P_CWP1_-FLuc* strain increased 400-fold when transferred to *in vitro* encystation medium (Supplemental Figure S2). This signal intensity was maintained for over three weeks of serial passage without antibiotic selection (Supplemental Figure S2B).

To determine the temporal and spatial dynamics of *Giardia* encystation *in vivo,* we inoculated eight mice with one million *P_CWP1_-FLuc* expressing trophozoites. *P_CWP1_-FLuc* bioluminescence was quantified every other day in live animals. One day post-infection we observed significant *P_CWP1_-FLuc* signal (Figure 3A), comparable to *in vitro* transcriptional upregulation of CWP1 (Supplemental Figure S2 and (23, 47)). The maximal BLI signal from the *P_CWP1_-FLuc* bioreporter occurred at 6 days post-infection, and significant signal ranged from five to eight days post inoculation (Figure 3B). While the *P_CWP1_-FLuc* bioluminescence from all animals was highest within the first week of infection, the bioluminescence was detectable throughout the 17 days of infection including day 1 (early infection), day 6 (mid-infection), and day 15 (late infection) (Figure 3B).

**Figure 3.**
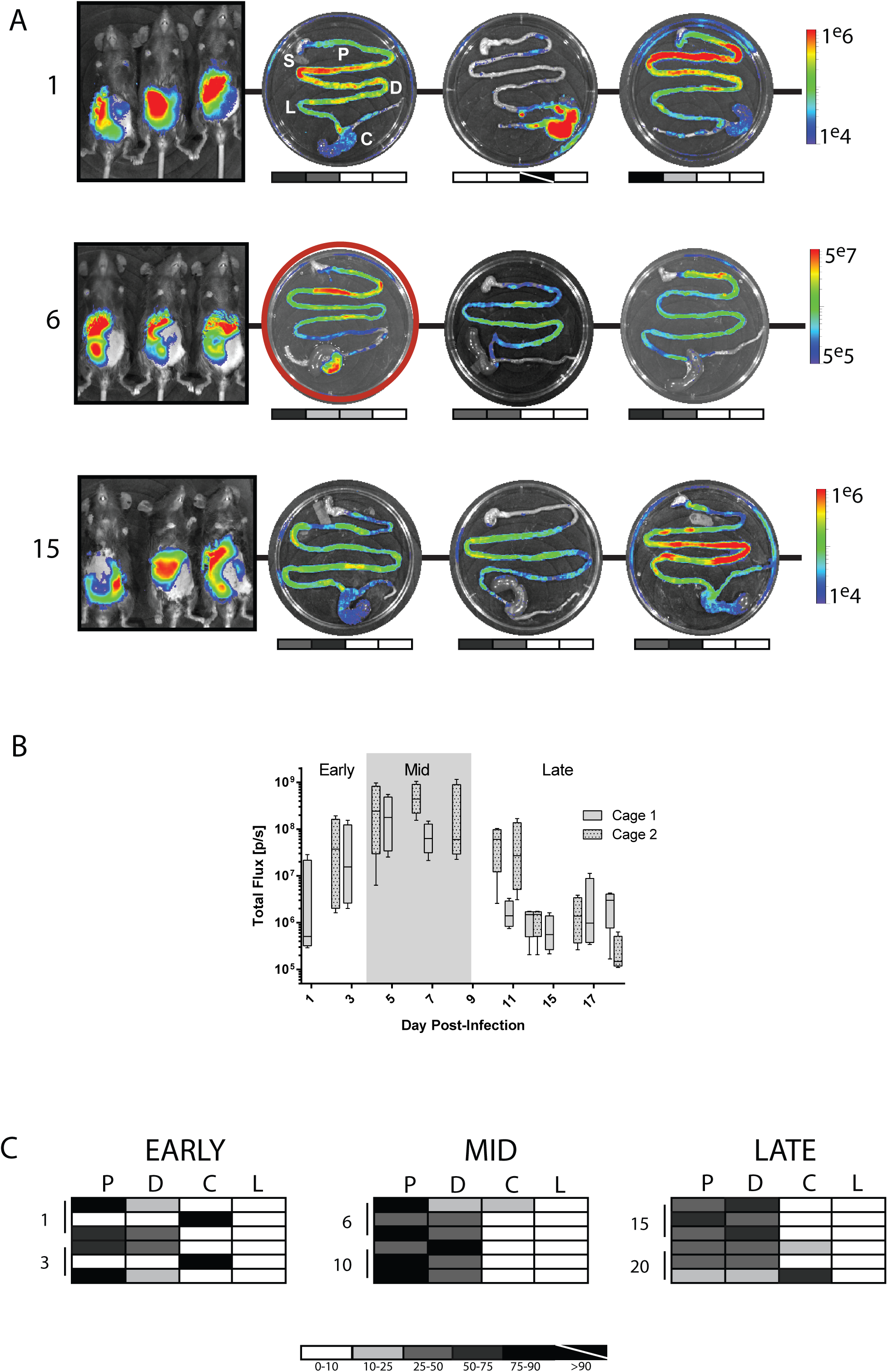
Encystation initiation occurs early in infection in both the proximal and distal small intestine. Eighteen mice were inoculated with *P_CWPT_-FLuc* strain, and cohorts of three animals were sacrificed and imaged on days 1, 3, 6,10,15, and 20 post-infection. In (A), the whole animal *in vivo* images from Days 1, 6, and 15, representing early, mid and late infection, are shown with corresponding *ex vivo* images from each animal (S = stomach, P = proximal, D = distal, C = cecum, and L = large intestine). Days 1 and 15 are presented on a scale between le^4^ and le^6^ photons/sec. Day 6 has the maximal signal, and images are presented using a scale between 5e^5^ and 5e^7^ photons/sec. For each *ex vivo* image, the photon flux (p/s/cm^2^/sr) for each intestinal segment is normalized to the maximal *ex vivo* bioluminescence signal on the radiance scale, yielding the percent total signal per segment. Gray scale maps of bioluminescence are shown below each *ex vivo* image (clear = 0-10% and black = 75-100%, with values between 10% and 75% indicated as shades of gray). In (B), two cages of mice **(N=4** per cage) were inoculated with the *P_CWP1_-FLuc* strain and imaged every other day. The box plot summarizes bioluminescent signal for encystation initiation *(P_CWPT_-FLuc)* for each phase of infection (**early,** days 0-3; **mid,** days **4-9;** and **late,** days 10-20). The bioluminescence from each cohort of animals is categorized by the gastrointestinal region, and the overall spatial localization of signal is summarized for each individual animal in each row of panel C. The shaded charts summarize the percentage of maximal bioluminescent signal from the *P_CWP1_-FLuc* strain in each of the four gastrointestinal regions (P = proximal small intestine, D = distal small intestine, C = cecum, and L = large intestine) for all infected animals in early, mid, and late stages of infection. The stomach (S = stomach) is shown for orientation but always lacks bioluminescence.

To determine the regions of the murine gut where encystation is initiated, cohorts of three animals were sacrificed on days 1, 3, 6,10,15, 20, and 26 post inoculation with the encystation bioreporter strain *P_CWPT_-FLuc*, and the entire Gl tract was imaged and scored by region (Figure 3A). Upregulation of the *P_CWPT_-FLuc* encystation bioreporter was detectable *ex vivo* as early as day one post infection. Maximal *P_CWPT_-FLuc* bioluminescence was primarily observed in the proximal small intestine, three to five centimeters distal to the stomach, similar to the constitutive *P_GDH_-FLuc* bioreporter strain. Like *P_GDH_-FLuc, P_CWP1_-FLuc* bioluminescence was often observed as regions of local maxima or foci within an area of lower bioluminescence (Figure 2 and Figure 3).

In contrast to the *P_GDH_-FLuc* bioreporter strain, bioluminescence from the encystation bioreporter *P_CWP1_-FLuc* was distributed throughout the entire small intestine (Figure 3A). Early in infection, equal numbers of mice displayed *P_CWPT_-FLuc* bioluminescence in the proximal and distal small intestines (Figure 3C), yet the bioluminescence from *P_CWP1_-FLuc* localizing to the proximal SI accounted for 50% of total intensity, whereas the distal SI signal was only 16% of the total bioluminescence in the gut. At mid and late infection, proximal and distal SI signal intensities were comparable (54% and 40%, peak; 37% and 44%, late) with equal numbers of mice showing signal from both proximal and distal SI regions.

We also noted increased localization of *P_CWP1_-FLuc* bioluminescent signal to the cecum, as compared to *P_GDH_-FLuc*, which localized primarily to the proximal small intestine (Figure 2). Two mice from early infection and one from late infection had strong cecal bioluminescent signals, sometimes at the exclusion of other anatomical sites, or in conjunction with bioluminescence elsewhere in the gastrointestinal tract.

### Confirmation of encystation initiation in the proximal small intestine during early infection

To confirm the encystation initiation pattern early in infection, we infected animals with a second encystation specific strain, *P_CWPT_-FLuc*, containing the promoter region of the cyst wall protein 2 (CWP2) gene (48). The temporal and spatial dynamics of encystation initiation we observed with *P_CWPT_-FLuc* were similar to those of *P_CWP1_-FLuc* (Supplemental Figure S6).

Because infections with both encystation-specific *P_CWP1_-FLuc* and *P_CWPT_-FLuc* bioreporter strains indicated that encystation initiation occurs early during infection and is primarily localized to the proximal SI, we confirmed the expression of CWP1 transcripts throughout the gut using qPCR of *ex vivo* samples following bioluminescent imaging. Within the first five centimeters of the proximal SI, transcription of CWP1 was upregulated by three days post infection, with significantly more upregulation by day 7 relative to basal CWP1 transcription levels in *in vitro* culture (Figure 4A).

**Figure 4.**
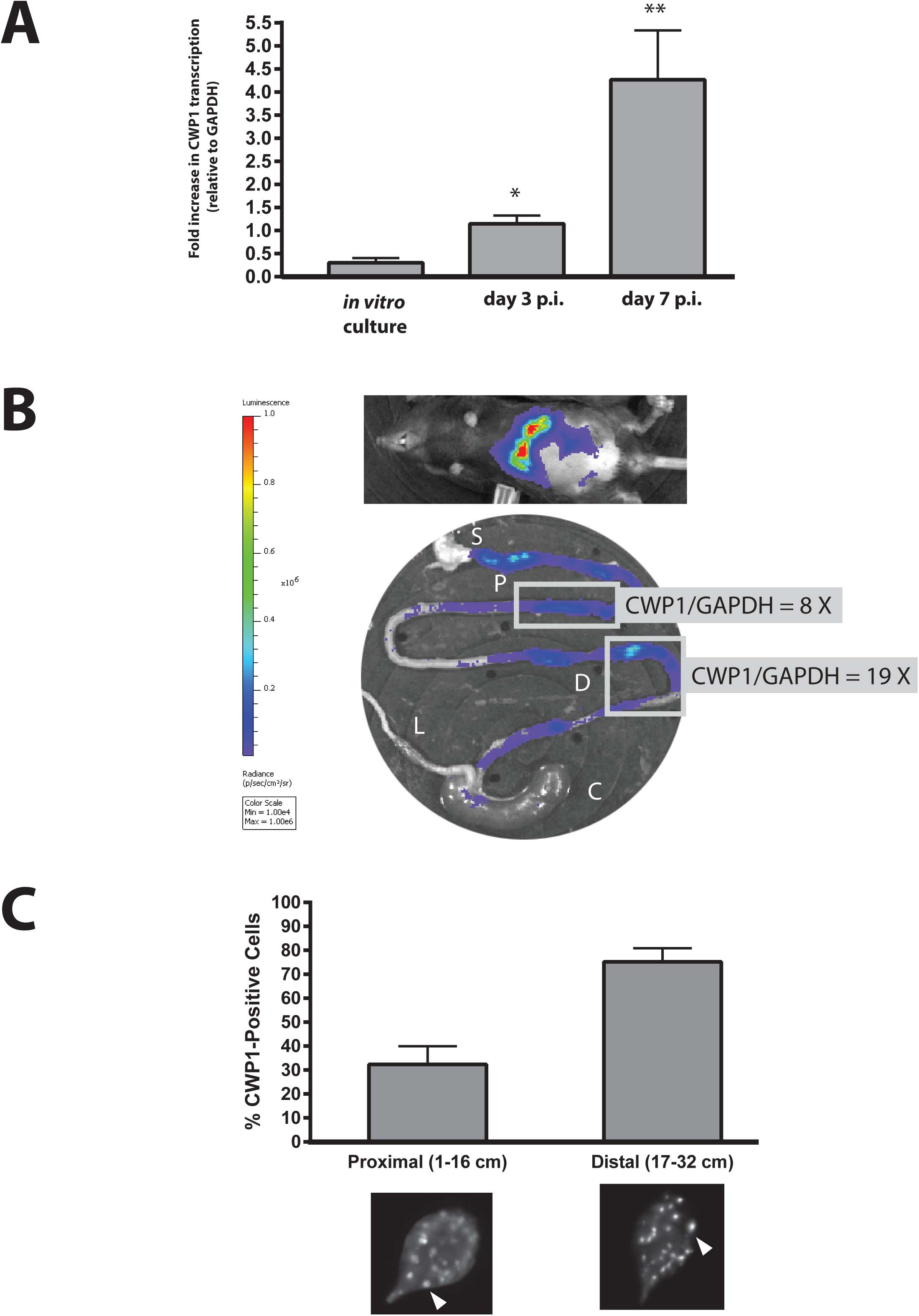
CWP1 qPCR and immunostaining of ESVs verify encystation in the proximal and distal small intestine. Quantitative PCR was used to compare the *in vivo* expression of CWP1 relative to GAPDH (CWP1/GAPDH) in the first five centimeters of the small intestine at day 3 and 7 p.i. to that of a confluent *in vitro* culture (A). Asterisks indicate statistical significance using unpaired T tests with Welch’s correction; one asterisk = p < 0.05 and two asterisks = p < 0.005. In B, a representative animal infected with the *P_CWPT_-FLuc* strain is imaged non-invasively at day 7 p.i., and *ex vivo* BLI of the gastrointestinal tract (P = proximal small intestine, D = distal small intestine, C = cecum, and L = large intestine) is shown with corresponding CWP1/GAPDH qPCR of representative proximal and distal small intestine regions (boxed). In C, encystation specific vesicles (ESV) were immunostained using an α-CWPl antibody and quantified in pooled *ex vivo* samples from two animals each at day 3 and 7 p.i. ESVs are marked by arrows in representative images from each respective intestinal sample. The percentages of trophozoites positive for ESVs are shown as a percentage of total trophozoites imaged (N>600 cells counted) in regions of the proximal small intestine (pSI) and distal small intestine (dSI).

Upregulation of encystation-specific promoter activity results in the commitment of trophozoites to differentiate into cysts that are shed into the environment to infect new hosts (47). Hallmarks of this commitment to encystation include the upregulation of CWP1 and CWP2 genes, and the appearance of encystation specific vesicles (ESVs) that transport the cyst wall proteins (e.g., CWP1 and CWP2) to build the cyst wall (49, 50). We find that *in vivo* CWP1 gene expression corresponds to the *in vivo* BLI signal of the *P_CWP1_-FLuc* strain. As previously shown in Figure 3, the *P_CWP1_-FLuc* bioluminescence is localized in foci throughout in the proximal and distal small intestine (Figure 4B). We observed a significant increase in CWP1 gene expression (relative to GAPDH) in these foci of the distal small intestinal and proximal small intestinal region. Specifically, we quantified 8 fold and 19 fold higher CWP1 gene expression in the proximal and distal small intestine, respectively, relative to GAPDH expression.

During *in vitro* encystation, ESVs appear within 5-7 hours following transfer to encystation medium (19, 47, 51). On days 3 and 7 post infection, we immunostained *ex vivo* intestinal samples using an anti-CWPl antibody (50) and confirmed that trophozoites with encystation-specific vesicles (ESVs) were also present throughout the small intestine (Figure 4C). CWP1 positive cells represented approximately 80% of the total cells imaged in the distal small intestine (Figure 4C). Specifically we identified at least twenty ESVs in each trophozoite positive fro ESVs that were isolated from the proximal regions of the small intestine (see representative images, Figure 4C).

### Increased parasite density contributes to encystation initiation

*Ex vivo* spatial imaging of bioluminescence showed that trophozoite colonization of the host gut is not uniform; rather, vegetative and encysting trophozoites are concentrated in foci, primarily within the proximal small intestine (Figure 2, Figure 3, and Supplemental Figure S6). Localized areas of increased parasite density might affect the physiology or differentiation of parasites in this particular region, perhaps contributing to developmental transitions. To assess whether the observed encystation promoter activity in mice was a consequence of initial concentrations of trophozoites used during gavage, we inoculated cohorts of mice (N=4 mice per group, n=12 total) with three different densities of *P_CWP1_-FLuc* trophozoites (Figure 5A). *P_CWPT_-FLuc* signal intensity was dependent on inoculum density during the first 6 days post-inoculation. After day 6, the bioluminescent signal reached a maximum that was similar for all three inoculum densities, with a slight and gradually decline over the next two weeks (Figure 5A). Of eight mice imaged daily for 14 days, the maximum bioluminescence was reached at an average of 6 days, with a range between five and eight days. With respect to the spatial location of encystation, at day 21 post infection, regardless of initial inoculation density, the *ex vivo* bioluminescent signal primarily remained in the proximal and distal small intestine (Figure 5B) although some animals had distal or cecum bioluminescence. We suggest that once the initial inocula reaches a colonization density threshold, perhaps localized to foci, encystation initiation occurs at the maximal level.

**Figure 5.**
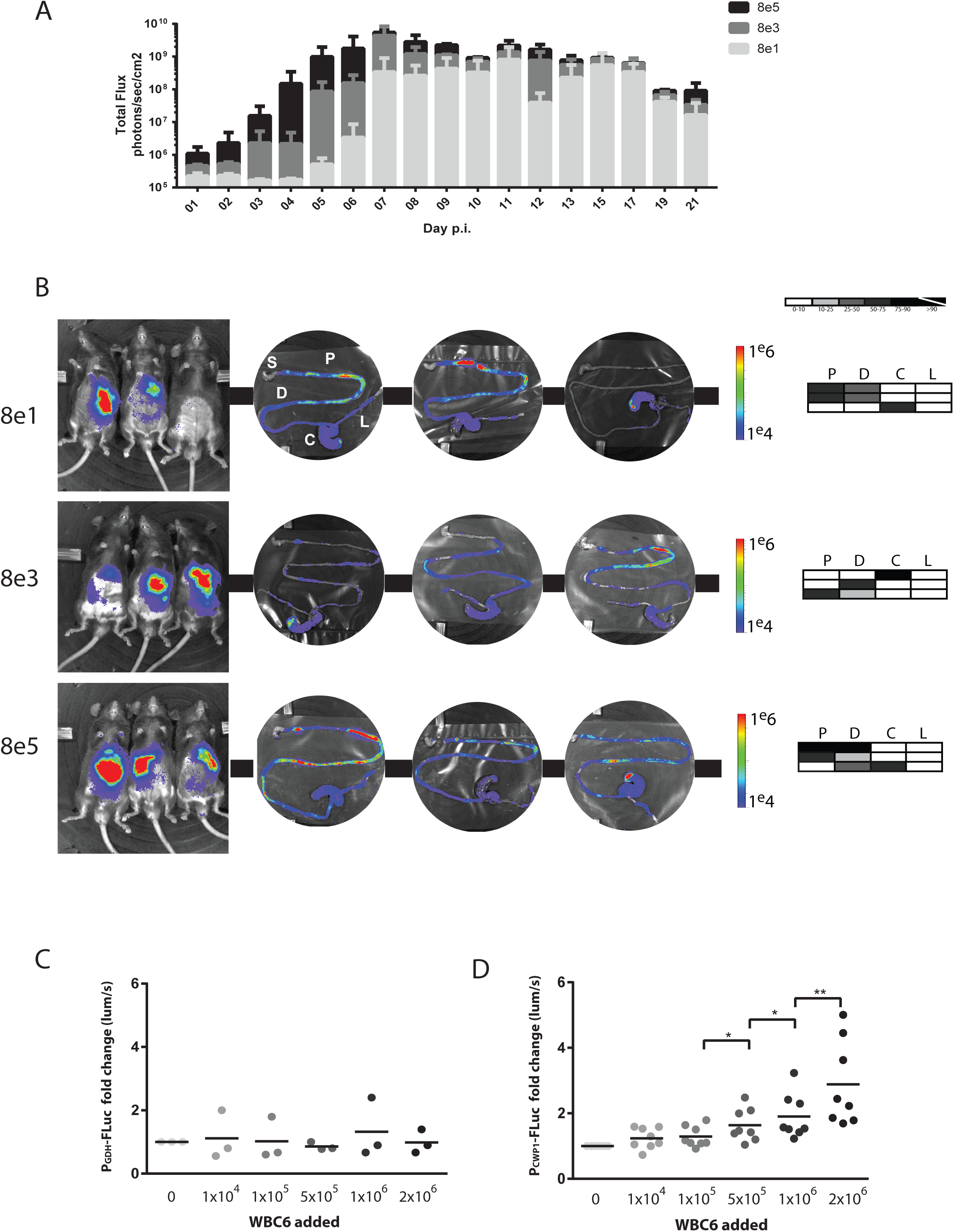
Increased cell density contributes to encystation initiation and upregulation of CWP1 expression. To assess the impact of cell density on the initiation of encystation, cohorts of three mice were infected with three different concentrations of *P_CWP1_-FLuc* parasites (80, 8000, or 800000) and bioluminescence was imaged and quantified daily over 21 days in total (A). In panel B, *in vivo* and *ex vivo* BLI is presented following sacrifice at day 21. The shaded charts in B summarize the percentage of maximal bioluminescent signal from the *P_CWPT_-FLuc* strain in each of the four gastrointestinal regions (P = proximal small intestine, D = distal small intestine, C = cecum, and L = large intestine) for each individual animal infected with that initial inoculum. The stomach (S = stomach) is shown for orientation but always lacks bioluminescence. In panels C and D, bioluminescence is quantified from an experiment in which 100,000 bioluminescent *P_GDH_-FLuc* (C) or *P_CWP1_-FLuc* (D) trophozoites were incubated up to twelve hours in encystation medium, and each well was crowded with increasing amounts (darker shaded dots) of non-bioluminescent WBC6 parasites (see Methods). The nine-hour time point is shown for both strains; additional time points are presented in Supplemental Figure *SI.* A single asterisk indicates p < 0.05 and a double asterisk indicates p < 0.01.

To evaluate whether parasite density had an effect on the initiation of encystation *in vitro,* we “crowded” cultures of the constitutive bioreporter *(P_GDH_-FLuc)* strain (Figure 5C) or the encystation-specific bioreporter *(P_CWPT_-FLuc)* strain (9 hours, Figure 5D) with increasing amounts of non-luminescent wild type WBC6 in encystation buffer. Then we quantified bioluminescence at 3, 6, 9 and 12 hours after transfer to encystation medium (9 hours, Figure 5C and 3-12 hours, Supplemental Figure S7). Within nine hours, we observed a significant increase in bioluminescence from the *P_CWP1_-FLuc* strain with the addition of 5×10^5^ −2×10^6^ additional trophozoites to the *P_GDH_-FLuc* (5D) as compared to no increase in bioluminescence with crowding of the *P_GDH_-FLuc* strain (Figure 5C).

## DISCUSSION

Despite its global burden, zoonotic potential, and emerging evidence of resistance to the most commonly used drugs (52, 53), *Giardia’s* pathogenesis in the host remains poorly understood. This dearth of information about the diversity of host-parasite interactions occurring within distinct regions of the gastrointestinal tract is primarily due to the lack of methods to directly and non-invasively interrogate disease progression in live animal hosts. Current standard methods of quantifying giardiasis involve sacrificing animals in cohorts at specific time points and determining parasite load by direct parasite counting or by G/’ord/o-specific qPCR (4, 54). Here we have developed and used non-invasive live bioluminescence imaging of giardiasis in mice to define *in vivo* spatiotemporal patterns of parasitic colonization and differentiation.

### Non-invasive quantification of *in vivo* temporal dynamics of giardiasis

Unraveling the complex relationship between host and parasite requires use of an *in vivo* model of infection. The most widely used animal model of giardiasis is infection of mice with human disease-associated *Giardia* strains (WB or GS) (28). Parasite burden in mice has been most commonly quantified by directly counting trophozoites in short segments of duodenum or jejunum, or more recently, by *Giardia*-specific qPCR (4, 54). Directly counting trophozoites in 2-3 cm segments of small intestine is laborious and subject to sampling errors. In live animals, quantification of fecal cysts is commonly used to estimate parasite abundance, yet cyst shedding may be a poor surrogate for overall parasite burden (55). Overall, bioluminescence imaging provides a faster and more sensitive evaluation of parasite burden than the current gold standard for parasite enumeration (56).

Non-invasive *in vivo* BLI relies on the external detection of light produced internally, and signal intensity may be limited by the overall level of luciferase expression, the oxygen tension within relevant tissues, pigmentation of organs and skin, or any background signal from the animal (35). However, the gut is sufficiently oxygenated to permit signal detection, and while animal tissues exhibit relatively high background levels of autofluorescence, they have nearly nonexistent levels of autoluminescence, which facilitates detection even at low signal strength (35, 37, 57). We monitored and quantified *in vivo* expression of metabolic and encystation genes for up to three weeks using cohorts of mice infected with one of three *Giardia* bioluminescent reporter strains *(P_GDH_-FLuc, P_CWP1_-FLuc* or *P_CWPT_-FLuc)*. Using the constitutively expressed *P_GDH_-FLuc* strain, we confirmed maximal infection at approximately seven days, consistent with prior studies of giardiasis in mice (28). This live imaging strategy provides the first real-time visualization of the spatial and temporal dynamics of giardiasis *in vivo,* allowing us to assess the timing and location of parasite differentiation. Cysts are known to be shed sporadically and sometimes cyclically (58, 59). Similar to the variability in human giardiasis (60), we also observed variability in infections between isogenic cage mates, including variations in the time to maximum infection, spatial colonization patterns, and cyclical infections (Figures 1–3). We confirmed that infection dynamics change based on the number of parasites inoculated, but that as a whole and as previously reported, parasite burdens peaks at about one week after infection.

### Reassessing the spatial dynamics of giardiasis in the mouse model

The convoluted route of the animal gut and the diffusion and refraction of bioluminescence presents a challenge when imaging giardiasis in live animals (34). Differentiating localized parasite activity from diffuse infection using *in vivo* optical imaging is also challenging (Fig 1 and 2). To resolve the spatial dynamics of *in vivo* infection, we used *ex vivo* imaging of bioreporters of both metabolic activity and encystation. Based on early studies using direct counting of trophozoites from intestinal samples, *Giardia* has generally been assumed to primarily colonize the mid-jejunum, or middle section of the small intestine (17, 61, 62), although other early work suggested that trophozoites prefer to colonize throughout the proximal small intestine (17). Rather than uniformly colonizing throughout a region of the Gl tract, we show that *Giardia* colonizes the intestine with a non-uniform or patchy distribution, characterized by discrete foci of colonization (Figure 2), while encystation initiation also occurs in discrete foci within the proximal and distal small intestine, or occasionally the cecum (Figure 3 and Supplemental Figure 5).

These high-density foci of infection are also upregulated in encystation initiation. Using *ex vivo* BLI, we imaged discrete foci of encysting trophozoites in the proximal small intestine and not other anatomical sites such as the cecum or large intestine. These observations challenge conventional assumptions that chemical cues in the distal gut are solely responsible for the initiation of *in vivo* trophozoite differentiation to the cyst in the *Giardia* life cycle. We found that metabolically active trophozoites are predominantly located in the proximal small intestine, with areas of local intensity frequently just distal to the pylorus (Figures 2 and 3). We also observed spatial variability between individuals, from diffuse infection throughout the small intestine to patchy foci only in the distal small intestine or cecum. Maximal bioluminescence (and thus infection) correlated strongly with proximal small intestinal colonization, whereas developing or clearing infections were present more diffusely throughout the gastrointestinal tract (Figure 3).

### High-density foci of parasites contribute to encystation initiation

The prevailing view of trophozoite differentiation to cysts in the host has been extrapolated from the chemistry of the region gut where trophozoites were believed to encyst (16-18). Parasite commitment to encystation and excystation are key events in *Giardia’*s life cycle, and it is clear that these transitions are highly regulated (47). Premature or tardy encystation can limit cyst development, proliferation, transmission into the environment, and dissemination to new hosts (55). The transition to the cyst form begins with detection of encystation stimuli, resulting in transcriptional upregulation of encystation-specific cyst wall proteins (CWPs) (23, 46). Almost 30 years ago, Gillin et al. showed that elevated bile concentrations, perhaps reminiscent of the distal small intestinal environment, could induce encystation in *in vitro* culture (20, 61). When these *in vitro* encystation protocols are used, CWPs are transported to the outer membrane via encystation-specific vesicles (ESVs) within roughly six hours of exposure to encystation stimuli (23, 46). However, differentiation to cysts can be induced even in the absence of bile, and several *in vitro* culture protocols have been developed to induce encystation by modifying pH, bile, lactic acid, and lipid concentrations in culture (18, 23). Importantly, no *in vitro* encystation protocol produces a high abundance of infectious cysts, suggesting that *in vivo* encystation may not accurately recapitulate differentiation *in vivo.*

Our quantitative *in vivo* imaging of temporal and spatial dynamics of parasite proliferation and encystation implies that there could be other factors contributing to *Giardia’*s developmental transitions during its life cycle in the host. Rather than encystation uniformly occurring throughout a particular region of the gut, we observed non-uniform foci of bioluminescence in infections with CWP1 and CWP2 strains (Figure 3 and Supplemental Figure 5). We also detected significant expression of CWP1 in *ex vivo* samples associated with increased bioluminescence in both the proximal and distal small intestine (Figure 4B). Lastly, we confirmed the presence of ESVs in trophozoites isolated from the proximal and distal small intestine (Figure 4), indicating that encystation is initiated and proceeds normally in these regions. In contrast with prior studies, this initiation of encystation occurred early in infections in the proximal small intestine (Figure 3) and peaked within the same time as maximal parasite density observed using the constitutively expressing bioreporter strain (Figure 1). While there was initial variation in the encystation bioluminescent signal proportional to the amount of initial inoculum, we saw that the encystation-specific BLI signal peaked about seven days and encystation BLI signal persisted throughout the twenty-one days of infection for all inoculum densities (Figure4A).

Pathogens have evolved to take advantage of the discrete mucosal surfaces and functions associated with the various anatomical regions of the mammalian gut (63). Reaching a particular threshold of cell density is known to either directly or indirectly modulate developmental programs in diverse parasitic (64-66) and free-living eukaryotes (67). Density-dependent quorum sensing is key to slender to stumpy differentiation in trypanosomes, for example, and trypanosomes can also respond to or affect bacterial quorum sensing signals (65, 66). Parasites such as *Giardia* detect and respond to a variety of chemical and environmental cues during their life cycles, and *Giardia* has been shown to respond to alterations in lipid and pH concentrations *in vitro,* triggering encystation.

We provide evidence that supports a model of a parasite density-based threshold for the induction of encystation in the proximal small intestine and in *in vitro* culture (Figure 5). In both our *in vivo* model of infection and *in vitro* culture experiments, the local parasite density of *Giardia* trophozoites may indirectly induce encystation. Compared to regions colonized with a lower parasite density, intestinal regions with higher parasite colonization density (foci), could directly impact the local chemistry of the gut, while indirectly impacting parasite physiology, the local gut microbiome, or the host epithelium (Figure 2). With respect to encystation initiation, foci of high parasite density could affect local concentrations of nutrients, metabolites, and pH — all of which are reported stimuli for encystation initiation (47). Thus we suggest that the nonuniform foci of encystation-specific bioluminescence (Figure 3) represent “hotspots” of encystation in the gut. Regardless of the initial number of trophozoites in the inoculum (Figure 5A) or the time post-infection (Figure 3), once the local density of trophozoites surpasses a specific threshold, encystation could be initiated. This model of localized parasite density-induced encystation due to the depletion of nutrients or accumulation of waste products is not incongruent with observed *in vitro* contributions of pH and/or lipid starvation to encystation initiation (16-18).

Throughout the gut, high density foci of parasites could create local changes in gut chemistry that could influence parasite metabolism and differentiation. In *Giardia,* the encystation signal could be induced either indirectly via localized nutrient deprivation and/or accumulation of waste products, or directly via an unrecognized quorum sensing-type mechanism for evaluating local cell density. Once trophozoites are committed to differentiation (47) in these localized foci within the gut, later developmental stages of encystation could occur more distally in the small intestine due to passage of encysting trophozoites through the gastrointestinal tract, resulting in the observed accumulation and/or shedding of cysts in the lower gut. While we observe some initiation of encystation within the first day of infection (Figure 3), the overall process of encystation in the host is lengthy, and it may take hours before mature, infectious cysts are observed in the large intestine or are recovered in feces. Further characterization of parasite physiology and differentiation in high density foci as compared to low density regions of colonization will help to elucidate the contribution of parasite density to *Giardia*’s developmental transitions.

### A new tool for evaluating chronic giardiasis and anti-giardial drug screening

Human giardiasis typically resolves within a few weeks, yet chronic or variable infections can occur (33) and have been linked to impaired physical and cognitive development in children (7). Chronic animal giardiasis models have not been possible because longitudinal analyses of infection dynamics in the same animal have not previously been feasible. *In vivo* BLI offers real-time and longitudinal monitoring of the infection dynamics of giardiasis, providing an animal model for long-term monitoring of giardiasis in cohorts of infected animals. In addition, animal numbers can be reduced with longitudinal BLI of giardiasis - a primary goal of ethical animal use in research (35, 68). Dual or triple spectra bioreporter strains (69-71) could be used to simultaneously assay two or more cellular processes (e.g., metabolic activity and encystation), integrating different components of infection into a single study animal over time.

Growing evidence of drug resistance in *Giardia* underscores the need to develop new therapeutic alternatives for the treatment of giardiasis (33), and *in vivo* bioluminescence imaging of murine giardiasis will aid in the evaluation of promising anti-giardial drug candidates. As we have shown, BLI of luciferase-expressing strains not only facilitates monitoring of parasite burden, but can also provide real-time information on other aspects of parasite physiology and metabolism. We have validated the use of BLI for the analysis of anti-*Giardia* drugs by demonstrating that metronidazole, the standard of care anti*-Giardia* drug that targets parasite metabolic activity (72), reduced *in vivo* bioluminescence of the constitutively-expressing *P_GDH_-FLuc* bioreporter strain. Other bioluminescent reporter strains could be utilized for high-throughput *in vitro* screens of candidate drugs, prior to *in vivo* assessment in mice. BLI imaging studies with anti*-Giardia* drugs targeting non-metabolic parasitic cellular processes (e.g., motility or encystation) could identify adjunct or complementary treatments that reduce parasite proliferation, infection duration, or cyst dissemination.

## Materials and Methods

### Luciferase strain construction and vaiidation

We created two strains of *Giardia iambiia* WBC6, each with firefly luciferase (FLuc) driven by a specific *Giardia* gene promoter (Supplemental Figure 1). FLuc promoter fusion constructs were integrated into the genome as previously described (38). To quantify colonization and metabolic activity, we integrated a construct containing FLuc driven by the constitutive NADP-specific glutamate dehydrogenase (GiardiaDB GL50803_21942) promoter *(P_GDH_-FLuc)* (Figure S1A). To quantify *in vivo* encystation dynamics, we integrated a construct containing FLuc with the encystation-specific cyst wall protein 1 (GiardiaDB GL50803_5638) promoter *(P_CWP1_-FLuc)* and the encystation-specific cyst wall protein 2 (GiardiaDB GL50803_5435) promoter *(P_CWP2_-FLuc)* (Figure SIB and SIC). Briefly, a vector previously used to integrate HA-tagged aurora kinase (73) was modified to contain the coding sequence for firefly luciferase fused to the GDH, CWP1, or CWP2 promoters. Puromycin (PurR) and ampicillin (AmpR) resistance cassettes allowed selection in *Giardia* and *E. coli,* respectively. The vector was linearized using *Miu*I and 10 |ig of DNA was electroporated into *Giardia lamblia* strain WBC6 (38). Transfected cells were selected for seven to ten days using puromycin (50 µg/ml). Confirmation of successful genomic integration was obtained by PCR amplification (data not shown), as well as *in vitro* bioluminescence assays in vegetative cells *(P_GDH_-FLuc)* and encysting strains *(P_CWP1_-FLuc* and *P_CWP2_-FLuc*) (Supplemental Figure S2).

### Giardia trophozoite and encystation culture conditions

*G. lamblia* (ATCC 50803) WBC6, *P_GDH_-FLuc, P_CWP1_-FLuc*, and *P_CWP2_-FLuc* strains were cultured in modified TYI-S- 33 medium supplemented with bovine bile and 5% adult and 5% fetal bovine serum [56] in sterile 16 ml screw-capped disposable tubes (BD Falcon), and incubated upright at 37°C without shaking. Encystation was induced *in vitro* by decanting TYI-S-33 medium from 24 hour cultures (roughly 30% confluent) and replacing with encystation medium modified by the addition of 0.5 grams/liter bovine bile, pH 7.8 (47). After 24 hours, cysts settled at the bottom of the tube.

### Giardia in vitro bioluminescence and density dependence assay

To assess the stability of luciferase signal in integrated promoter-FLuc strains without selection, luciferase expression in the *P_GDH_-FLuc* and *P_CWP1_-FLuc* strains was determined before and after passage of the cells in the absence of antibiotic selection (1:25 dilutions daily for three weeks). Confluent tubes were incubated on ice for 15 minutes to fully detach cells. Cells were pelleted by centrifugation at 900 × g for 5 minutes and resuspended in 1 mL of fresh TYI-S-33 media supplemented with 150 µg/mL D-Luciferin (PerkinElmer). Aliquots (50 µl, three technical replicates) were added to white opaque 96-well microplates (Perkin Elmer). Bioluminescence was analyzed on a Victor3 plate reader using one-second exposures until maximum signal was achieved.

For density dependence assays, wild-type and *P_CWP1_-FLuc* cells were grown to confluence, harvested as described, and washed and resuspended in encystation media. One hundred thousand *P_CWPT_-FLuc* cells were plated in each well of a microplate and a range of dilutions of non-bioluminescent wild-type WBC6 were added to the *P_CWP1_-FLuc* cells in three technical replicates. Encystation medium was then added to adjust the final volume to 200 µL per well. Microplates were individually sealed in Type A Bio-Bags (BD) to maintain an anoxic environment and incubated at 37°C for the indicated time points. D-luciferin was added to 150 µg/mL and luciferase activity was analyzed as described previously.

### Non-invasive in vivo bioluminescent imaging of Giardia colonization and encystation

Eight week old, female C57/B6/J mice (Jackson Laboratories) were maintained on *ad libitum* water and alfalfa-free irradiated rodent pellets (Teklad 2918). To promote parasite colonization, water was supplemented with 1 mg/ml ampicillin and neomycin (Teknova) for five days prior to infection (54). Water bottles were kept protected from light to minimize degradation of the antibiotics and antibiotics were refreshed every other day. Individual mice were marked with ear tags or permanent marker on tails, and hair was removed from the ventral abdomen to facilitate imaging. Each mouse was first shaved using a commercial men’s groomer, and then depilatory cream (Nair) was applied for 30 seconds. For long-term studies, depilatory cream was re-applied as necessary to maintain a hairless ventrum (34). Each animal was gavaged with 1 × 10^7^ *G. lamblia* trophozoites in 100 µL phosphate-buffered saline as previously described (74). All animal studies were performed with IACUC approval at the University of California Davis (Scott C. Dawson, PI).

For *in vivo* BLI, mice were sedated using isoflurane (1.5-3%) in an induction chamber. D-luciferin (30 mg/kg) was then injected intraperitoneally at a dose of 150 mg/kg body weight (total volume injected, 100 µL). Sedated mice were transferred to an optically clear XIC-3 Isolation Chamber (PerkinElmer) and positioned on their dorsal surface. Bioluminescence was imaged using an IVIS Spectrum (PerkinElmer) with no emission filter. Anesthesia was maintained with 1.5-2% isoflurane and 100% oxygen during imaging.

Photons were quantified using an ultra-sensitive CCD camera (IVIS Spectrum) and the resulting heat maps of bioluminescent photon emission intensity were overlaid on still images of anaesthetized animals. To allow the D-luciferin to distribute throughout the body, images were collected with two-minute exposures constantly over 8-10 minutes until the bioluminescent signal stabilized. The final image collection was performed with two to five minute exposures, dependent on signal strength. ROI analysis was used to quantify bioluminescence (LivingImage). A rectangle encompassing the entire abdomen was drawn for each mouse from front paws to anus. BLI data was quantified as total flux (photons/second) for exposure time-independent quantification of signal intensity. For mice infected with *P_GDH_-FLuc,* the minimal signal was normalized to the level of background signal in uninfected mice (lxlO^4^ photons/sec). Because the bioluminescent signal intensity from *P_CWPT_-FLuc* was several orders of magnitude stronger than for mice infected with for *P_GDH_-FLuc,* the minimal threshold signal was adjusted to 5x10^5^ photons/sec in order to minimize background.

### Ex vivo bioluminescence imaging

Sedated mice were euthanized by cervical dislocation. The gastrointestinal tract was quickly dissected from esophagus to anus and positioned within a plastic Petri dish. The dish and contents were placed within the XIC-3 Isolation Chamber, 2.5% oxygen was provided to maximize signal, and the Gl tract was imaged with a two-minute exposure. *Ex vivo* imaging was performed less than 30 minutes after the initial injection of luciferin. ROI analysis was used to quantify bioluminescence (Livinglmage). Total gastrointestinal tract signal was analyzed with a circle over the entirety of the petri dish. The stomach, proximal SI (first half), distal SI (second half), cecum, and large intestine were traced using the free-hand tool.

### Estimation of parasite density using quantitative PCR (qPCR)

One-centimeter segments from a region showing strong *ex vivo* signal were identified, marked in the Livinglmage software, excised, and flash frozen in liquid nitrogen. Total genomic DNA was extracted using standard methods (75), and diluted to 10 ng/µL in nuclease-free water prior to qPCR. Quantitative PCR of the pyruvate-ferredoxin oxidoreductase-1 *(pfor1,* GiardiaDB GL50803_17063) gene (72) was performed using PforlF 5′TTCCTCGAAGATCAAGTTCCGCGT3′ and PforlR 5′TGCCCTGGGTGAACTGAAGAGA AT3′ oligonucleotide primers and SensiFast No-ROX cyber-green master mix in an MJ Opticon thermal cycler, with an initial 2 minute denaturation step at 95°C followed by 40 cycles of 95 °C for 5 seconds, 60°C for 10, and 72 °C for 10. The single copy, constitutively expressed murine nidogen-1 (*nid1*) gene was used as an internal control to quantify the contribution of murine DNA to intestinal segments (qPCR primers nidoF 5′CCAGCCACAGAATACCATCC3′, nidoR 5′GGACATACTCTGCTGCCATC3′). The differential counts to threshold (ΔCT) between *nid1* and *pfor1* were quantified, and Ct values were determined using the Opticon Monitor software.

### Confirmation of encystation in the proximal small intestine during early infection using quantitative PCR (qPCR)

A cohort of four mice was infected with *P_CWP1_-FLuc* and two mice were imaged and sacrificed for sample collection at both day 3 and day 7 p.i. Following *ex vivo* imaging, intestines from infected mice were dissected into 3 cm segments, immediately frozen in liquid nitrogen and transferred to a −80°C freezer until used for RNA extraction. RNA from intestinal segments was purified using RNA Stat-60 (Tel-test, Inc.). RNA quality was assessed using spectrophotometric analysis (NanoDrop Technologies) and electrophoresis prior to cDNA synthesis. Double-stranded cDNA was synthesized using the QuantiTect Reverse Transcription Kit (Qiagen). Quantitative PCR of cyst wall protein 1 (CWP1, GiardiaDB GL50803_5638) was performed using cwplF 5′TAGGCTGCTTCCCACTTTTGAG 3′ and cwplR 5′AGGTGGAGCTCCTTGAGAAATTG 3′oligonucleotide primers (76)and SensiFast No-ROX cyber-green master mix in an MJ Opticon thermal cycler, with an initial two minute denaturation step at 95°C followed by 40 cycles of 95 °C for 5 seconds, 60°C for 10 s, and 72 °C for 10 s. The constitutively expressed gene for glyceraldehyde 3-phosphate dehydrogenase (GAPDH, GiardiaDB GL50803_ 6687) was chosen as an internal reference gene and was amplified with gapdh-F 5′ CCCTTCACGGACTGTGAGTA 3′ and gapdh-R 5′ ATCTCCTCGGGCTTCATAGA 3′ oligonucleotide primers. Ct values were determined using the Opticon Monitor software and statistical analyses were conducted using Prism (GraphPad).

### Immunostaining of encysting trophozoites in intestinal tissue samples

Intestinal segments from the same infected mice analyzed with qPCR (above) were transferred to chilled *Giardia* medium, incubated at 4°C for 15 minutes to allow parasite detachment, then briefly vortexed. Cells remaining in the supernatant were pelleted at 900 × g for 2 minutes at 4°C, and resuspended in 0.5 ml of chilled medium. Parasites were then spread on pre-warmed (37°C) glass coverslips and incubated 30 minutes in a humidifying chamber to facilitate attachment. Two ml of warmed fixation buffer (4% paraformaldehyde, 1% glutaraldehyde in 0.86X HEPES buffered saline) was added to each coverslip and incubated at room temperature for 15 minutes. Coverslips were washed three times with 2 ml of PEM buffer, and then incubated in quenching buffer (0.11 M glycine in 0.9X PEM) for 15 minutes at room temperature. Immunostaining was performed as previously described (77) with a mouse primary antibody against cyst wall protein 1 (Giardi-a-Glo, Waterborne, Inc.) and a donkey anti-mouse secondary antibody conjugated to an Alexa 350 fluorophore (Invitrogen).

Images were acquired via automated Metamorph image acquisition software (MDS Technologies) using a Leica DMI 6000 wide-field inverted fluorescence microscope with a PlanApo 100X, NA 1.40 oil immersion objective. At least 100 trophozoites were counted per slide and cells were binned into encysting or normal trophozoite morphologies. Regions of small intestine were distinguished spatially as follows: proximal 1-4 cm, proximal 5-16 cm, and distal 17-32 cm. Slides from 3-6 separate intestinal segments were counted per spatial bin. The statistical significance of differences in cell number between the spatial bins was determined via Student’s T-Test.

## ACKNOWLEDGEMENTS

Bioluminescent imaging was performed at the Center for Molecular and Genomic Imaging (CMGI), University of California, Davis. We would like to acknowledge Jennifer Fung and Charles Smith for help with training and acquisition of images. The authors graciously thank Kari Hagen and Hannah Starcevich for critical reading of the manuscript.

